# Detection and accurate False Discovery Rate control of differentially methylated regions from Whole Genome Bisulfite Sequencing

**DOI:** 10.1101/183210

**Authors:** Keegan D. Korthauer, Sutirtha Chakraborty, Yuval Benjamini, Rafael A. Irizarry

## Abstract

With recent advances in sequencing technology, it is now feasible to measure DNA methylation at tens of millions of sites across the entire genome. In most applications, biologists are interested in detecting differentially methylated regions, composed of multiple sites with differing methylation levels among populations. However, current computational approaches for detecting such regions do not provide accurate statistical inference. A major challenge in reporting uncertainty is that a genome-wide scan is involved in detecting these regions, which needs to be accounted for. A further challenge is that sample sizes are limited due to the costs associated with the technology. We have developed a new approach that overcomes these challenges and assesses uncertainty for differentially methylated regions in a rigorous manner. Region-level statistics are obtained by fitting a generalized least squares (GLS) regression model with a nested autoregressive correlated error structure for the effect of interest on transformed methylation proportions. We develop an inferential approach, based on a pooled null distribution, that can be implemented even when as few as two samples per population are available. Here we demonstrate the advantages of our method using both experimental data and Monte Carlo simulation. We find that the new method improves the specificity and sensitivity of list of regions and accurately controls the False Discovery Rate (FDR).

## 1. Introduction

DNA methylation is an important epigenetic modification that plays a role in a wide variety of biological processes. Numerous studies have been carried out to locate CpG loci where DNA methylation may be involved in gene regulation, differentiation, and cancer. With recent advances in sequencing technology such as Whole Genome Bisulfite Sequencing (WGBS), it is now possible to measure DNA methylation at single base resolution across all CpGs in the genome. Even though the most common application of the technology is to detect differentially methylated regions (DMRs) between populations, most methods for analysis of WGBS experiments focus on statistical differences for CpG loci one at a time (Akalin *et al.*, 2012, Dolzhenko and Smith, 2014, Lee and Morris, 2016, Park *et al.*, 2014, Park and Wu, 2016). While useful, approaches for identification of differentially methylated loci (DML) have many practical limitations in both implementation and interpretation. Here, we discuss these limitations as well as outline the challenges of performing inference at the region level. Finally, we introduce a rigorous statistical approach that overcomes these challenges to construct de novo DMRs with accurate FDR control.

Methods to identify DMLs in WGBS experiments are greatly hindered by the high-dimensionality and low sample size setting that is common in high-throughput genomics studies. The number of tests performed is equal to the number of loci analyzed, which is very large in typical WGBS studies. In the human genome, for example, there are close to thirty million CpG loci (Smith and Meissner, 2013). Further, DML methods generally do not account for the well-known fact that measurements are spatially correlated across the genome (Leek *et al.*, 2010) and instead treat measurements from all loci as independent. Correcting for multiple comparisons without taking into account these correlations can result in a loss of power.

Additionally, methods for assessing the significance of DMLs typically require large sample sizes due to reliance on large sample approximations (Dolzhenko and Smith, 2014, Hansen, Langmead and Irizarry, 2012, Hebestreit, Dugas and Klein, 2013, Lee and Morris, 2016). Although WGBS is the current gold standard for estimating whole genome methylation profiles (Marx, 2016), cost limitations are still a barrier to acquiring more than a few individuals per biological condition in many studies (Ziller *et al.*, 2015). This is reflected in the study design of major consortiums that aim to characterize the epigenome. For example, WGBS experiments in murine embryos carried out as part of the ENCODE project are limited to two biological replicates per tissue type and developmental time point combination (He *et al.*, 2017). In addition, the number of biological replicates measured with WGBS in the UCSD Human Reference Epigenome Mapping Project (Schultz *et al.*, 2015) is also limited to 2-3 per tissue type. As such, we aim to maximize power while controlling the false discovery rate even with sample sizes as small as two samples per condition.

Methods for identifying DMLs also need to properly model count data that does not conform to standard Gaussian models. This is in contrast to methylation array analysis, where Gaussian models performed well (Jaffe *et al.*, 2012). One option is to assume that methylation proportions, defined as the number of methylated reads divided by the number of total reads covering a given CpG locus, follow a normal distribution (Hansen, Langmead and Irizarry, 2012). However this assumption clearly does not hold when the total reads covering the CpG, referred to as the coverage, is small, a common occurrence in these datasets. The approach also ignores that variance of this proportion depends on the coverage. To overcome these limitations, DML approaches have also modeled WGBS count data using Binomial models (Saito, Tsuji and Mituyama, 2014). However, Binomial models on their own cannot account for biological variability within sample groups. In order to account for biological variability in count data, Beta-Binomial models (Park *et al.*, 2014, Sun *et al.*, 2014) are a natural extension. However they come at the cost of increased computational burden when testing millions of loci.

Beyond implementation challenges, DML approaches also suffer from limited interpretability. In general, identifying DMRs is more biologically relevant than reporting DMLs. Apart from the so-called ‘CpG traffic lights’ (Khamis *et al.*, 2017), most individual CpG loci likely do not have a large impact on epigenetic function on their own, but rather through a biochemical modification that involves several loci. Most notably, regional DNA methylation levels are correlated with the expression levels of nearby genes. Specifically, methylation gain is associated with stable transcriptional silencing of nearby genes (Bird, 2002). In the context of differential methylation analysis, Aryee *et al.* (2014) found that differentially expressed genes were consistently more likely to be located near DMRs than DMLs.

While DML approaches may construct DMRs by chaining together neighboring significant loci, this type of approach will not yield a proper assessment of the statistical significance of the constructed regions, nor will the False Discovery Rate (FDR) be properly controlled (Robinson *et al.*, 2014). This is because controlling the FDR at the level of individual loci is not the same as controlling FDR of regions, as has been noted in the context of peak calling in ChIP-seq experiments (Lun and Smyth, 2014, Siegmund, Zhang and Yakir, 2011). FDR correction at the level of individual loci means that the proportion of expected false positive loci is controlled, not the proportion of false positive regions. Statistically, this is a critical point since FDR control of DMR detection is not guaranteed under the DML setting. In fact, many discoveries at the loci level may constitute only a single discovery. This means that a large number of correct rejections at the loci level can inflate the denominator in the FDR calculation, which will artificially lower the false discovery rate of loci as compared to regions (Figure 1). We were motivated to develop a procedure to control FDR at the region level and provide an accurate measure of statistical significance for each region.

**Figure 1:**
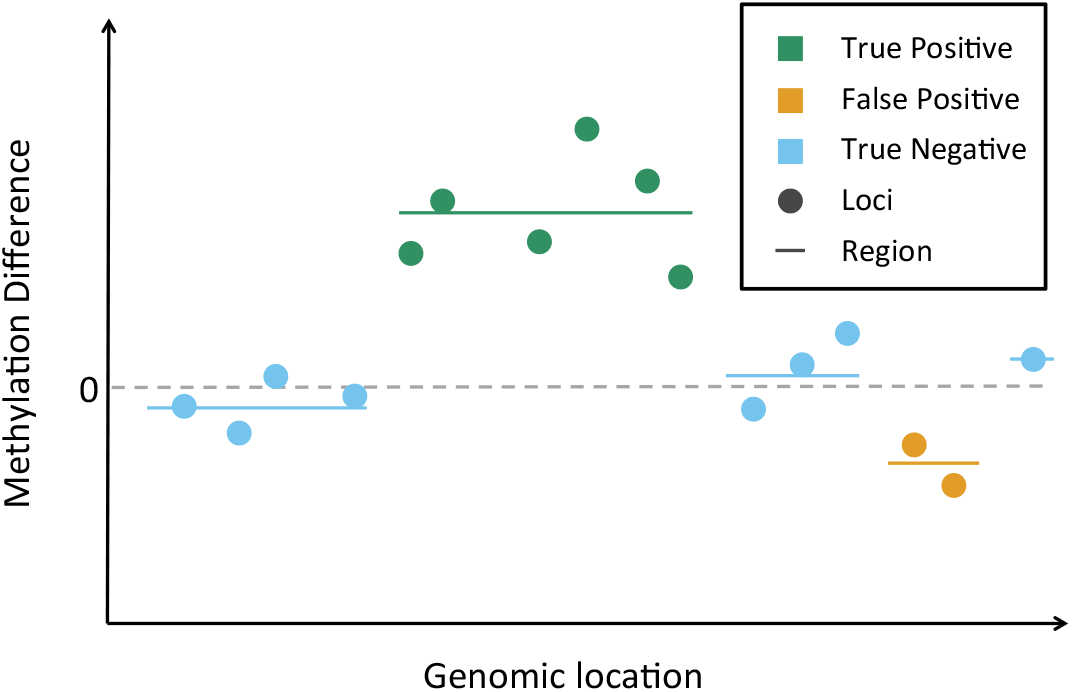
Illustration of why FDR at the loci level is not the same as FDR at the region level. This schematic shows a plot of genomic location versus methylation difference estimates at several neighboring loci. The individual CpGs (points) are shaded by whether they are a true or false positive. Regions are denoted by lines. The loci 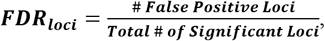 which is equal to 0.25 in this example. The region 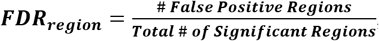 which is equal to 0.50 in this example.

Many recent computational approaches have been developed with the goal of identifying DMRs, but most do not provide formal inference for regions (Hansen, Langmead and Irizarry, 2012, Saito, Tsuji and Mituyama, 2014, Wu *et al.*, 2015, Yu and Sun, 2016) and instead join together significant DMLs. This type of procedure will suffer from the problems outlined above. Other approaches can perform inference at the region level, but only for predefined regions of interest or fixed sliding windows (Hebestreit, Dugas and Klein, 2013, Sun *et al.*, 2014). Though useful in targeted settings such as Reduced Representation Bisulfite Sequencing (RRBS), or when we have prior knowledge of the DMR size, they are not applicable to identifying DMRs of arbitrary size from WGBS. Those methods that scan the genome for DMRs and provide inference at the region level do not properly control FDR (Juhling *et al.*, 2016, Wen *et al.*, 2016). This is evidenced, for example, by the FDRs reported in the simulation studies of Wen *et al.* (2016), which were as high as 0.85 and widely varied across scenarios. Juhling *et al.* (2016) also do not achieve accurate FDR control in simulation studies (see Section 4.1).

The challenge of performing inference at the region level is complicated by several factors in addition to the challenges already discussed in the context of DML analysis. The first challenge is in defining the region boundaries themselves. Without prior knowledge or predefined regions, we need to construct data-driven regions. Calculating a test statistic for these data-driven regions of varying sizes with a known null distribution is not straightforward. In addition, challenges are presented by the complex statistical dependencies observed in measurements from nearby loci (Benjamini, Taylor and Irizarry, 2016), as well as different within group variability across loci (Hansen, Langmead and Irizarry, 2012). Some methods ignore correlation across loci (Wen *et al.*, 2016) or biological variability from sample to sample (Saito, Tsuji and Mituyama, 2014, Wu *et al.*, 2015). Not properly accounting for both of these sources of variability in DNA methylation data, however, results in misleading conclusions or loss of power. For a full review of DML and DMR methods, see (Shafi *et al.* 2017).

To overcome the limitations and challenges detailed above, we propose a two-stage approach that first detects candidate regions and then explicitly evaluates statistical significance at the region level while accounting for known sources of variability. Candidate DMRs are defined by segmenting the genome into groups of CpGs that show consistent evidence of differential methylation. Because the methylation levels of neighboring CpGs are highly correlated, we first smooth the signal to combat loss of power due to low coverage as done by Hansen, Langmead and Irizarry (2012). In the second stage, we compute a statistic for each candidate DMR that takes into account variability between biological replicates and spatial correlation among neighboring loci. Significance of each region is assessed via a permutation procedure which uses a pooled null distribution that can be generated from as few as two biological replicates, and false discovery rate is controlled using the procedure of Benjamini and Hochberg (1995). Code to reproduce the analyses presented in this paper is provided in Supplementary material and the open-source R package dmrseq that implements the approach is available on GitHub.

In Section 2, we provide a detailed description of the datasets used. We describe the methodological details of the approach and detail the data processing and analysis procedure in Section 3. In Section 4, we present our findings using both experimental data and simulations. We demonstrate that the proposed approach assigns greater statistical significance to regions that have greater biological significance in terms of potential functional roles in the regulation of gene expression. We also evaluate sensitivity and specificity of the approach by analyzing null comparisons of samples from the same biological condition, with and without adding simulated DMRs. We demonstrate that dmrseq has higher sensitivity than existing approaches and accurately assesses statistical significance of regions through False Discovery Rate estimation. A discussion of the advantages and limitations of the method are given in Section 5.

## 2. Data Description

dmrseq is generally applicable to WGBS data which contains the counts for both methylated and unmethylated reads mapping to each CpG loci. This information can be obtained from raw sequencing reads using the mapping software Bismark (Krueger and Andrews, 2011), as described in the Supplementary materials. Specifically, CpG loci that are covered by at least one read in every sample should be used in the analysis. Other methods for analysis of WGBS data recommend removing CpG sites that have only a few reads in each sample, and while processed data of this form may be analyzed by our approach, it is important to note that this may result in a loss of power to detect regions in low-coverage areas of the genome.

In this study, we use our approach to identify DMRs using publically available WGBS data from two different case studies, as described below. We also evaluate sensitivity and specificity of DMR methods by applying them to simulated data. Summary of coverage and methylation values for all datasets used can be found in Table 1 and Supplementary Figure S2. For more details on data processing, see Section 1 of the Supplementary materials.

**Table 1:**
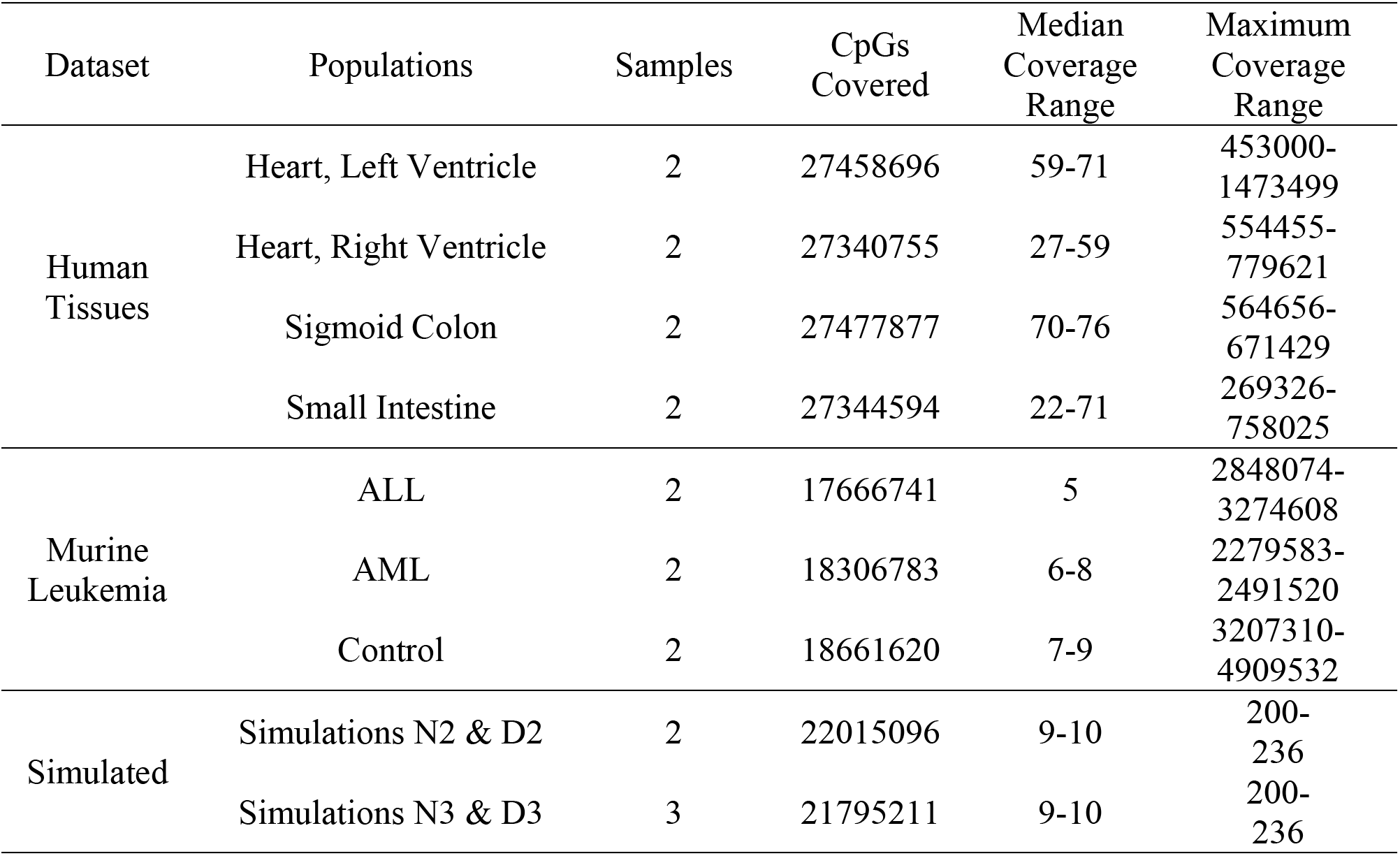
Summary of datasets used. Summary measures include the number of samples per population (‘Samples’), the number of CpGs with at least one read in all samples in the population (‘CpGs Covered’), median number of reads mapping to each covered CpG (‘Median Coverage’), minimum and maximum number of reads mapping to each covered CpG (‘Coverage Range’). Since the number of CpGs and their coverage are identical in the null comparisons and DMR simulations, the entries for N2 and D2 are combined. Likewise for N3 and D3.

### 2.1 Simulated data

Two sets of simulated data were constructed: one representing a null comparison (with no DMRs) and another containing simulated DMRs. To ensure that the simulated datasets closely match the characteristics of the observed experimental data, they were generated based on WGBS data from a study of human dendritic cells (Pacis *et al.*, 2015). This study estimated methylation profiles of human dendritic cells from six donors before and after infection with a pathogen. The null comparison was constructed by randomly partitioning the six control samples (before infection) into two groups of three samples each, denoted Simulation N3. The same is done for a subset of four of the samples to evaluate performance when there are only two samples in each population, denoted Simulation N2.

Starting with the null comparisons, 3,000 simulated DMRs were added to each dataset in order to evaluate specificity and sensitivity. These are denoted Simulations D2 and D3 for two and three samples per population, respectively. Briefly, a DMR is constructed by sampling a cluster of neighboring CpGs and simulating the number of methylated reads, conditional on observed coverage, for the samples from one population from a binomial distribution. The binomial probabilities are equal to the observed methylation proportions plus or minus a randomly sampled difference, which varies smoothly over the region according to a function similar to the tricube kernel (Cleveland, 1979) (see Section 2.4 of the Supplementary materials).

### 2.2 UCSD Human Reference Epigenome Mapping Project

Data from several human tissue samples from the UCSD Human Reference Epigenome Mapping Project (Schultz *et al.*, 2015) was used to identify DMRs related to tissue type. Specifically, four tissues were selected for performing pairwise comparisons: (1) Heart, left ventricle, (2) Heart, right ventricle, (3) Sigmoid colon, and (4) Small intestine.

### 2.3 Murine models of leukemia

In this study, marrow or thymus cells from two biological replicates form each of three different murine lines were extracted and genome-wide methylation levels measured with WGBS. One condition consisted of a wild-type control mouse. The other two had alterations in one or both of the DNMT3a or FLT3 loci, both of which have previously demonstrated implications in the development of leukemia (Pacis *et al.*, 2015). The mouse model with a wild-type DNMT3a locus and a duplication of the FLT3 locus has been shown to induce ALL. The mouse model with the same duplication of the FLT3 locus as well as a knock out of DNMT3a has been shown to induce the more lethal and aggressive AML. The DNMT3a also plays a role in promoting DNA methylation, so it is of interest to characterize the resulting differences in methylated regions among the control and two different leukemia models.

## 3. Analysis Framework

A two-step procedure is carried out to (1) construct de novo candidate regions, and (2) score candidate regions to quantify the effect of the covariate of interest on methylation level, and evaluate statistical significance by comparing them to null regions. Here we detail each stage of the approach.

### 3.1 Construction of candidate regions

In step 1, we detect candidate regions that contain multiple loci showing evidence of a difference in the smoothed pooled methylation proportion between biological conditions. For simplicity of presentation, we assume there are two biological conditions *s* ∈ {1,2}, with sample indices *j* ∈ *C_s_* (see Supplementary materials Section 2.7 for the case of more than two conditions). Let *M_ij_* be the number of methylated reads and *U_ij_* the number of unmethylated reads for locus *i* of sample *j* from condition *s*. The coverage is denoted *N_ij_*, where *N_ij_* = *M_ij_* + *U_ij_*. The estimate of the mean methylation proportion 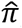 for loci *i* in condition *s* is taken to be the sum of methylated reads from all samples in that condition divided by the sum of all reads (i.e. the coverage) from all samples in condition *s*:

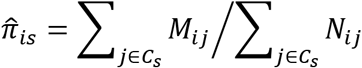

This leads to the following estimate of methylation proportion difference between 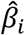 condition *s* and *s′* at loci *i*:

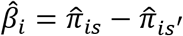

In order to give more weight to measurements with higher coverage, this estimate pools together samples within the same condition. To account for biological variability between samples and further reduce influence of observations with low coverage, smoothed individual loci estimates 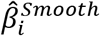 are obtained using a local-likelihood smoother (Loader, 1999) with smoothing weights *w_i_* equal to the median coverage at loci *i* scaled by the average Median Absolute Deviation (MAD) within the sample groups 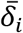:

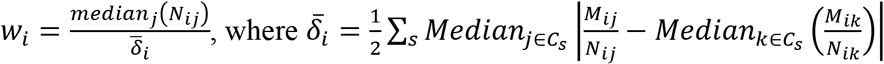

This places more emphasis on observations with high coverage and low variability within sample group (see Section 2.1 of the Supplementary material for more details).

Candidate regions are defined by segmenting the genome into groups of loci with a smoothed and scaled pooled proportion difference 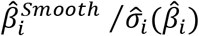 in the same direction that is greater than some threshold in absolute value (refer to Supplementary materials Section 2.2 for more details). Maximum spacing between loci within a candidate region is controlled by a predetermined value, and loci at the start and end of the region with low difference values are trimmed (refer to Supplementary materials Section 2.3 for more details). The threshold value should be chosen liberally so that it will more or less capture all of the true differences without regard to false positives, as significance of the candidate regions is assessed in the next step.

### 3.2 Assessing significance of regions

In the second step, we assess the significance of candidate regions. This task is complicated by the fact that the null statistics are calculated on an enriched set of regions. In general, the null distribution generated by the type of selection procedure described in the previous section is not known. A natural approach would be to carry out a permutation test to control FWER (family-wise error rate), which is done by Jaffe *et al.* (2012) to infer DMRs from array data. However, this is not feasible when we have only a few samples per population as is most often the case with WGBS. Thus, we set out to construct a statistic that can be comparable across the genome so that the signal can be compared among regions. Such an exchangeable statistic allows us to generate an approximate null distribution by pooling genomewide candidate regions detected from permutations.

To generate an approximately exchangeable region statistic that measures the strength of methylation difference, we need to account for sources of variation that are known to vary across the genome, including biological variability from sample to sample (Hansen, Langmead and Irizarry, 2012), as well as covariance of nearby loci (Benjamini, Taylor and Irizarry, 2016). Failing to do so may result in large test statistics just by chance for regions with high variability, leading to increased FDR or decreased power. For example, if we use an area-based statistic (Hansen, Langmead and Irizarry, 2012) or a mean difference statistic averaged across loci, power to detect DMRs is greatly reduced in simulation studies (Supplementary Figure S5 and Supplementary materials Section 4.1).

Since we need to compute the statistic over potentially hundreds of thousands of candidate regions, we also favor an approach that provides efficient and stable estimation procedures. For these reasons, we make use of generalized least squares (GLS) regression model with a nested autoregressive correlated error structure for the effect of interest on transformed methylation proportions, the advantages of which are described in detail in the next subsections.

#### 3.2.1 Estimation of region statistics with Generalized Least Squares models

To account for sampling variability, we assume that methylation counts for region *r* are Binomially-distributed with probability *p_ijr_*, where

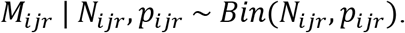

To model biological variability, we allow the binomial proportion for samples in condition *s* ∈ {1,2} to vary according to a beta distribution with shape parameters *α_irs_* and *β_irs_*, where

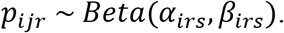

Let 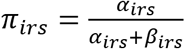 denote the mean of this Beta distribution. We are interested in estimating and assessing the significance of the difference in mean methylation levels across a region *r* for two biological conditions.

Our approach models transformed methylation proportions using GLS to obtain an approximation of the effect of interest. While directly modeling counts with either a Beta-Binomial Generalized Linear Model (GLM) or a Generalized Linear Mixed Model (GLMM) would allow us to accommodate complex covariance structures across samples and loci, it also results in complex likelihoods that require iterative maximization for each candidate region. Further, these procedures are subject to instability of estimation for methylation levels near the boundaries (zero and one) or non-identifiability in the case of separation as they occur in GLM (Gelman *et al.*, 2008) and GLMM (Abrahantes and Aerts, 2012) estimation. GLS models, in contrast, are efficient and stable to estimate due to the availability of approximate closed-form parameter estimates. Though GLS does not model counts directly, we incorporate information lost after transformation of methylation proportions through specification of a variance estimate that depends on coverage.

We choose the arcsine link function *Z_ijr_* = *arcin*(2*M_ijr_*/*N_ijr_* - 1) to obtain transformed methylation proportions, as proposed by (Park and Wu, 2016) for DML analysis, for its desirable ability to stabilize the dependence of the variance on the mean methylation level. While the variance of methylation proportions *M_ijr_*/*N_ijr_* depends on the mean parameter *π_ijr_*, the variance of *Z_ijr_* only depends on coverage *N_ijr_* and the dispersion of the Beta-Binomial distribution (refer to Supplementary materials Section 2.6 for more details). This helps us to form a statistic involving the transformed proportions that is exchangeable across regions that have different mean methylation values.

We assume a linear effect on the arcsine link-transformed methylation proportion parameters:

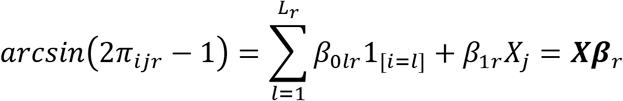

Here *β*_0*lr*_ are loci-specific intercept terms that account for variation on overall methylation levels across the region, where *l* = 1,…*L_r_* and *L_r_* denotes the number of loci in region. The coefficient for the effect of interest (e.g. biological group) is *β_1*r*_*. We denote the design matrix as ***X*** and the (*L_r_* + 1)-length vector of all coefficients 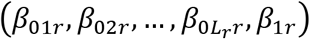 as ***β**_r_*. This leads to the following model for the transformed response 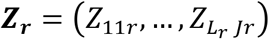 in region *r*

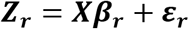

where we assume that *E*[ε_***r***_] = 0 and *Var*[ε_***r***_] = ***V_r_***, which can be fit by GLS given an estimate of the covariance matrix ***V_r_***. Since GLS allows arbitrary covariance structures, we use an autoregressive correlation structure to account for the correlation of methylation levels among nearby loci. To account for the dependence of the variance on coverage as mentioned above, we use variance weights. More details on the specific structure and estimation of ***V_r_*** are given in the next section.

With the above model, we assess the strength of the effect of the covariate of interest on methylation level within region *r* using the t-statistic *t_r_* from the Wald test of the null hypothesis that *β*_1*r*_ = 0. Parameter estimates and their standard errors are obtained with the ‘gls’ function in the ‘nlme’ package (Pinheiro *et al.*, 2017). Significance is evaluated by permutation using a pooled null distribution as described in detail in Section 3.2.3.

#### 3.2.2 Covariance of methylation levels within regions

In the estimation of the covariance matrix ***V_r_***, we take into account biological variability through variance weighting, and correlation of nearby loci through an autocorrelation structure. The variance weighting is done to account for the dependence of the variance of transformed values *Z_ijr_* on coverage. This variance depends non-linearly on *N_ijr_* (Supplement Section 2.6), but in order to enable efficient closed-form estimation with GLS, we further approximate it by

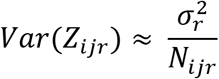

In addition, in order to construct a valid permutation test where the variance conditional on the effect of interest is invariant to permutation, we assume this variance identical for all samples at a given loci by approximating *N_ijr_* by *median_j_*(*N_ijr_*) = *N_i.r_*.

To model correlation of nearby loci, we use the flexible continuous autoregressive correlation structure of order 1, abbreviated CAR(1). Under CAR(1), the correlation parameter depends on the length of the interval between the two observations considered in the following manner

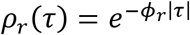

where *τ* is the length of the interval between two observations and *ϕ_r_* is the positive continuoustime autoregressive coefficient (following the notation of Jones and Boadi-Boateng (1991)) for region *r*. Thus, for subject *j*, the predicted methylation value for loci *i* at location *t_ijr_* in region *r* given the methylation value at loci *i* - 1 is

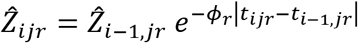

If the error variance of the CAR1 process is 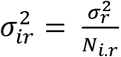 and we let the correlation structure be nested within subject (i.e. such that observations from two subjects are independent), it follows that the covariance matrix for a given sample can be written

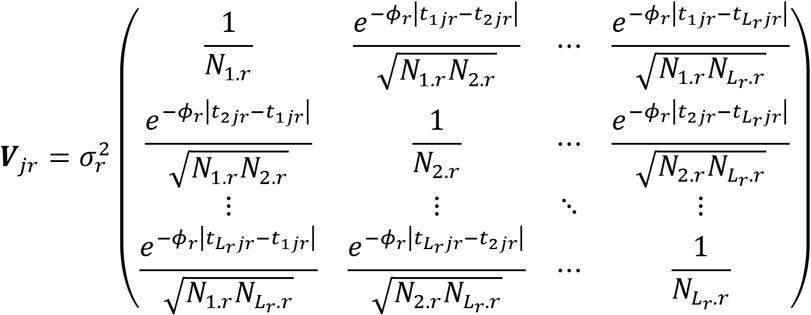

and for two subjects *j* and *j′*,*Cov(Z_ijr_,Z_ij′r_)* = 0.

The estimation of *ϕ_r_* is computationally efficient to carry out on small to moderately sized regions. However, for larger regions with more than 40 loci we use the slightly simpler AR(1) correlation structure since it is many times faster to compute. This discrete formulation assumes that observations are equally spaced, and that observations that are separated by lag 1 are correlated with region-specific correlation parameter *ρ_r_*. In addition, observations that are separated by *m* positions are correlated by 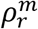. This results in a covariance matrix for ***Z_r_*** from region *r*, subject *j* of

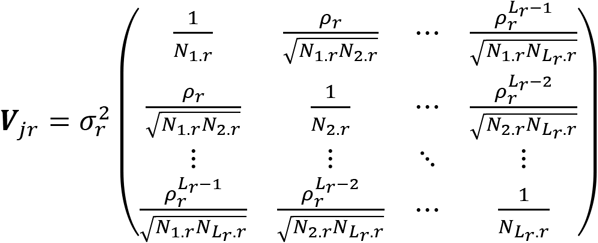

and for two subjects *j* and *j′*,*Cov(Z_ijr_,Z_ij′r_)* = 0.

The CAR(1) structure simplifies to the AR(1) process under certain conditions when observations are equally spaced (Jones and Boadi-Boateng, 1991). Thus the discrete AR(1) can be viewed as an approximation of the CAR(1) when correlations are positive and the two provide increasingly more similar estimates as observations approach constant spacing. Indeed, when comparing model fits under both correlation structures in simulated data, the t-statistics for the coefficient of interest under CAR(1) generally converge to the estimates under AR(1) as the number of loci increases (Supplementary Figure S1 and Section 2.5).

#### 3.2.3 Permutation to generate a null set of regions

The values of the covariate of interest (e.g. biological group) are permuted and the previous steps repeated in order to generate a set of statistics under the null hypothesis. Since the statistics account for known sources of variation that would otherwise prevent to comparison of regions across the genome, we can pool them together to form an approximate null distribution with as few as two samples per population. The empirical p-value is calculated by comparing the observed test statistics to the entire null set of statistics from all permutations. Control of FDR is carried out by adjusting the p-values using the procedure of Benjamini and Hochberg (1995).

## 4. Results

For each of the datasets described in Section 2, we applied dmrseq, as well as three widely used methods for DMR detection: BSmooth (Hansen, Langmead and Irizarry, 2012), DSS (Park and Wu, 2016), and metilene (Juhling *et al.*, 2016). Each approach was evaluated based on the criteria detailed in the next subsections. For specific details on software implementation, refer to the Supplementary materials (Section 3).

### 4.1 Simulation using dendritic cell data

Specificity was evaluated by identifying DMRs in null comparisons of two (N2) and three (N3) samples per group. Sensitivity was evaluated by identifying simulated DMRs in comparisons of two (D2) and three (D3) samples per group. Performance of each method is assessed by its ability to identify as many of the simulated DMRs as possible, while identifying as few DMRs as possible in the null comparison.

dmrseq did not identify any DMRs at the 0.05 level for the null comparisons N2 or N3 (Table 2). This remains true even when increasing the FDR threshold to 0.5 in both settings. In contrast, metilene identified a small number of DMRs, DSS identified many hundreds, and BSmooth tens of thousands using default settings (specific parameter specifications provided in Supplementary materials Section 2.6). When applied to the datasets with simulated DMRs (D2 and D3), dmrseq is able to accurately control the False Discovery Rate, whereas metilene cannot (Figure 2, Supplementary Figure S3). Note that analogous results cannot be obtained from DSS or BSmooth, as there is no way to specify FDR level.

**Table 2:** Null comparison results for sample size 2 (N2) and sample size 3 (N3). Numbers of DMRs identified by dmrseq and metilene are shown at the 0.05 FDR level. Default settings were used for BSmooth and DSS.

| Null Comparison | Method | DMRs (FPs) |
| --- | --- | --- |
| N2 | dmrseq | 0 |
|  | BSmooth | 76,563 |
|  | DSS | 661 |
|  | metilene | 31 |
| N3 | dmrseq | 0 |
|  | BSmooth | 76,319 |
|  | DSS | 770 |
|  | metilene | 27 |

**Figure 2:**
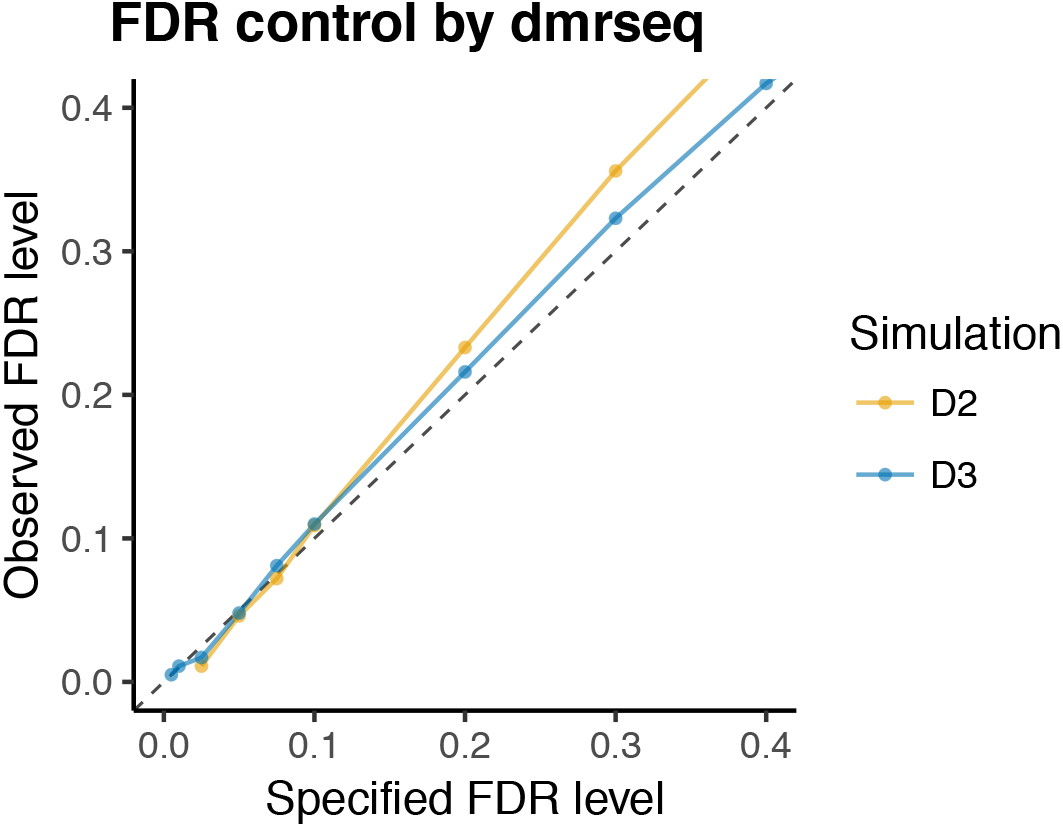
dmrseq provides accurate FDR control of regions. Specified versus observed region-level FDR level is plotted for two different sample size settings from simulated data for dmrseq. Note that region-level FDR cannot be specified for BSmooth or DSS, and results for metilene are shown in Supplementary Figure S3.

BSmooth and DSS identify similar numbers of False Positive regions in D2 and D3 compared to the null setting of N2 and N3, and far more than dmrseq and metilene (Table 3). Although both BSmooth and DSS have favorable numbers of TPs, it is clear that this comes at the expense of lack of control of FDR (Figure 3). Similarly, metilene has favorable numbers of FPs, but this comes at the expense of low power. Further, even at similar observed FDR levels, dmrseq achieves higher power levels than the alternative methods.

**Figure 3:**
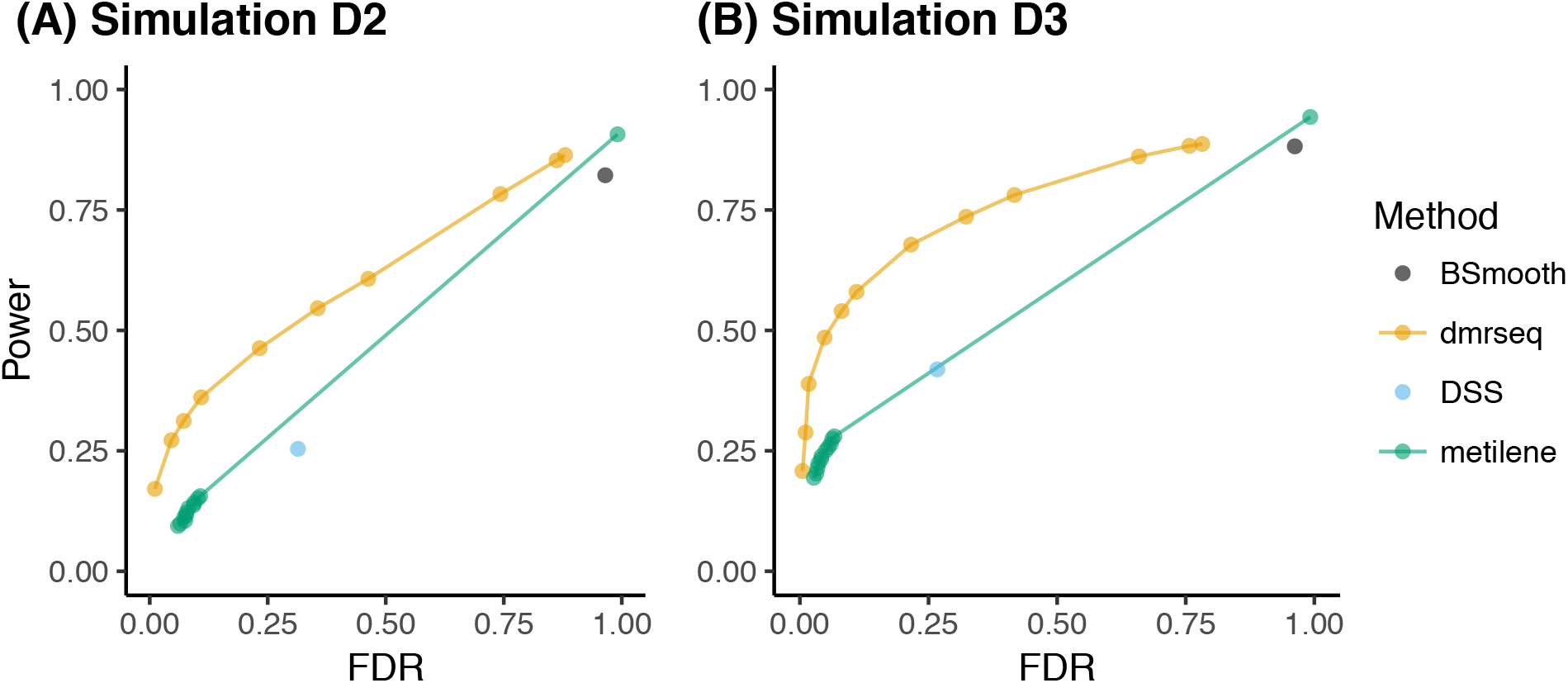
dmrseq is more powerful than other methods. FDR and power results for (A) Simulation D2 and (B) Simulation D3, with method denoted by color. dmrseq and metilene results are displayed for several different FDR cutoffs. Since region level FDR control is not possible for BSmooth and DSS, results using default settings are displayed. Power is calculated as the proportion of simulated DMRs overlapped by at least one identified DMR. FDR is calculated as the proportion of DMRs identified that do not overlap with any of the simulated DMRs.

**Table 3:** Simulated DMR results for sample size 2 (D2) and sample size 3 (D3). Numbers of DMRs identified by dmrseq and metilene are shown at the 0.05 FDR level. Default settings were used for BSmooth and DSS. True Positives (TPs) is the number of simulated DMRs that are overlapped by at least one identified DMR. False Positives (FPs) are DMRs that do not overlap any of the simulated DMRs.

| Simulation | Method | DMRs | TPs (unique) | FPs |
| --- | --- | --- | --- | --- |
| D2 | dmrseq | 914 | 816 | 42 |
|  | BSmooth | 73,252 | 2,466 | 70,688 |
|  | DSS | 2,086 | 762 | 655 |
|  | metilene | 329 | 210 | 30 |
| D3 | dmrseq | 1,620 | 1,455 | 78 |
|  | BSmooth | 72,764 | 2,646 | 69,999 |
|  | DSS | 2,858 | 1,257 | 763 |
|  | metilene | 652 | 441 | 27 |

Although FDR thresholds are not available for BSmooth or DSS, we also investigated the sensitivity and specificity of other settings beyond defaults of the thresholds at the single-loci level (the loci t-statistic cutoff for BSmooth, and the loci p-value for DSS). Making these thresholds more conservative generally reduced the numbers of False Positives, but once again dmrseq was consistently able to identify more True Positives at similar numbers of False Positives (See Supplementary Results and Figure S4).

We also stress that although lower False Positive rates could be achieved in this simulation study for BSmooth and DSS, individual loci thresholds do not correspond directly to specific FDRs at the region level. As a result, in practice, one must choose a threshold either by default settings, or by trial and error.

### 4.2 Human tissue and murine leukemia experimental data

The human tissue and murine leukemia studies were evaluated empirically based on the observed association of DMRs with differential expression by RNA-seq. Differentially expressed (DE) genes were identified using DESeq2 version 1.14.1 (Love, Huber and Anders, 2014). To assess functional relevance of the results, detected DMRs that overlap promoter regions of DE genes were assessed for signal in the expected direction. Specifically, a DMR - DE gene pair is expected to have higher methylation values in the sample group with lower expression. The odds that the DMR and DE statistics are in opposing directions are calculated at various FDR cutoffs for dmrseq and metilene to assess whether top-ranked DMRs are more likely to be biologically relevant. The same is done for various cutoffs for the numbers of top-ranking regions by effect size. Additionally, for each cutoff we calculate the number of CpGs covered and the proportion of detected DMRs that are within 2kb (from the center of the region) of a promoter region of a DE gene.

To qualitatively assess the ability of the dmrseq region-level summary statistic to rank DMRs as compared to other methods, we display example regions from the human tissue and murine leukemia studies. These examples illustrate the increased variability of regions that are highly ranked by naïve statistics but not dmrseq (Figure 5). We include a DMR with concordant rankings that exhibits clear differences between two human tissue types (Figure 5A). In contrast, the regions with discordant rankings between dmrseq q-value and mean difference (Figure 5B) and area statistics (Figure 5C) exhibit considerable variability between samples or loci (See Supplementary materials Section 2.8 for more details).

**Figure 5:**
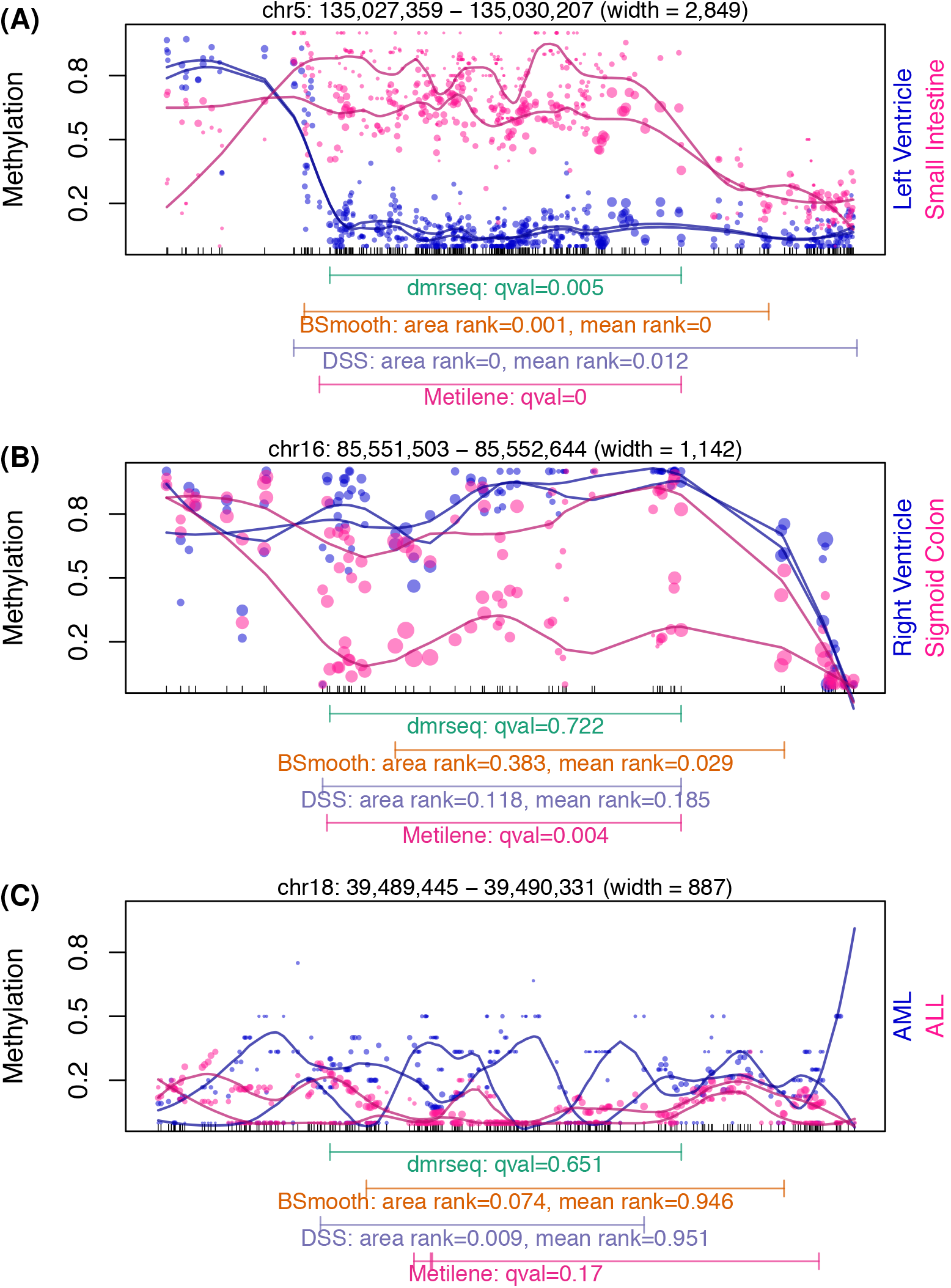
dmrseq ranks regions by statistical significance. Example regions from the human tissue and murine leukemia studies are displayed for three cases that illustrate the increased variability of regions that are highly ranked by area or mean difference statistics of BSmooth and DSS but not dmrseq. For each case, the q-value is shown for dmrseq and metiline, and the rank percentile by the area statistic and mean difference statistics are both shown for BSmooth and DSS (see Supplement Section 2.8 for details). (A) All methods assign a consistently high rank. (B) dmrseq assigns a low rank, but the mean difference statistic of BSmooth and DSS assign a high rank. (C) dmrseq assigns a low rank, but the area statistic of BSmooth and DSS assign a high rank. The condition comparison is indicated by the labels to the right of each plot.

#### 4.2.1 Tissue specificity in human samples

For DSS, metiline, and dmrseq, the number of DMRs found (Table 4) parallels the numbers of DE genes found by DESeq2 (Supplementary Table S2), but DSS generally found far more DMRs and metline far fewer. For BSmooth, however, the number of DMRs identified was similar for all comparisons. This happens because the cutoff for the individual loci statistics is set by default at a quantile of the observed statistics, resulting in a similar number of loci being deemed significant.

**Table 4:**
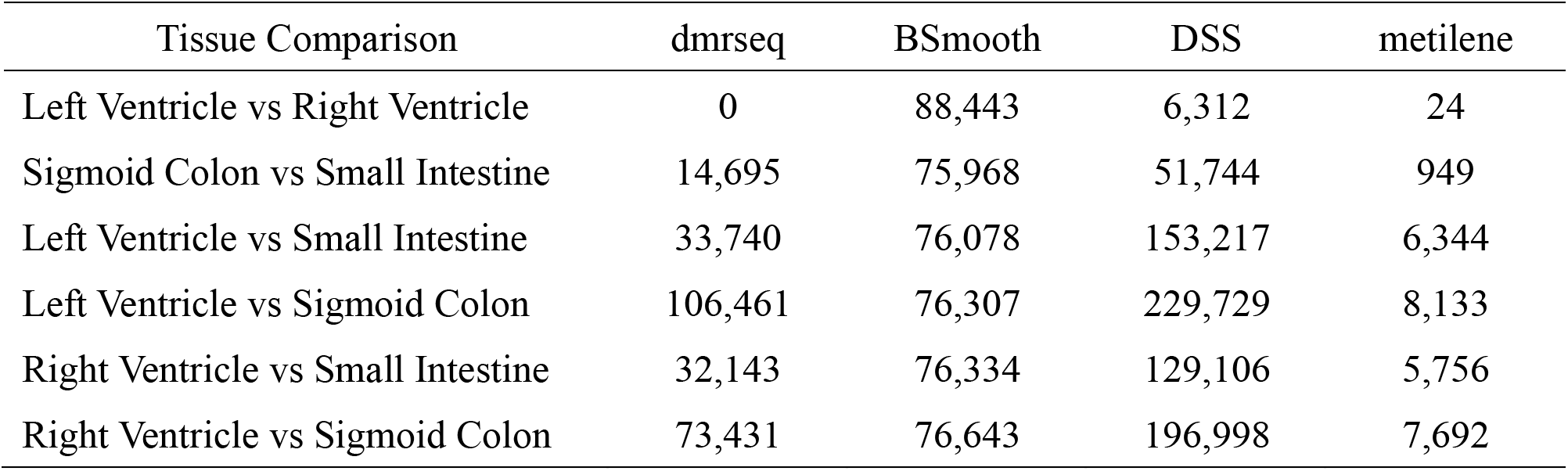
Tissue-specific DMR results. Number of DMRs found by dmrseq and metilene at FDR level 0.05, and BSmooth and DSS under default settings.

| Tissue Comparison | dmrseq | BSmooth | DSS | metilene |
| --- | --- | --- | --- | --- |
| Left Ventricle vs Right Ventricle | 0 | 88,443 | 6,312 | 24 |
| Sigmoid Colon vs Small Intestine | 14,695 | 75,968 | 51,744 | 949 |
| Left Ventricle vs Small Intestine | 33,740 | 76,078 | 153,217 | 6,344 |
| Left Ventricle vs Sigmoid Colon | 106,461 | 76,307 | 229,729 | 8,133 |
| Right Ventricle vs Small Intestine | 32,143 | 76,334 | 129,106 | 5,756 |
| Right Ventricle vs Sigmoid Colon | 73,431 | 76,643 | 196,998 | 7,692 |

The tissue-specific DMRs found by dmrseq are enriched for inverse associations with DE genes, and this enrichment is stronger for DMRs with lower FDRs (Figure 4). Additionally, enrichment of dmrseq DMRs is generally stronger than that of alternative methods. While metiline also provides an FDR estimate, there is no consistent association between the FDR ranking and strength of association with expression. DMRs identified by BSmooth and DSS cannot be ranked by FDR and the default settings may not be ideal, so we also rank DMRs by effect size (raw methylation difference) with optimized parameter settings (see Supplementary materials Section 3.2). The BSmooth and DSS DMRs with highest effect sizes exhibit comparable enrichment to dmrseq, with metilene considerably lower (Supplementary Figures S6 and S7). However, arbitrary cutoffs of effect size do not directly correspond to significance level, and the enrichment when including all DMRs is highest for dmrseq (Figure 4).

**Figure 4:**
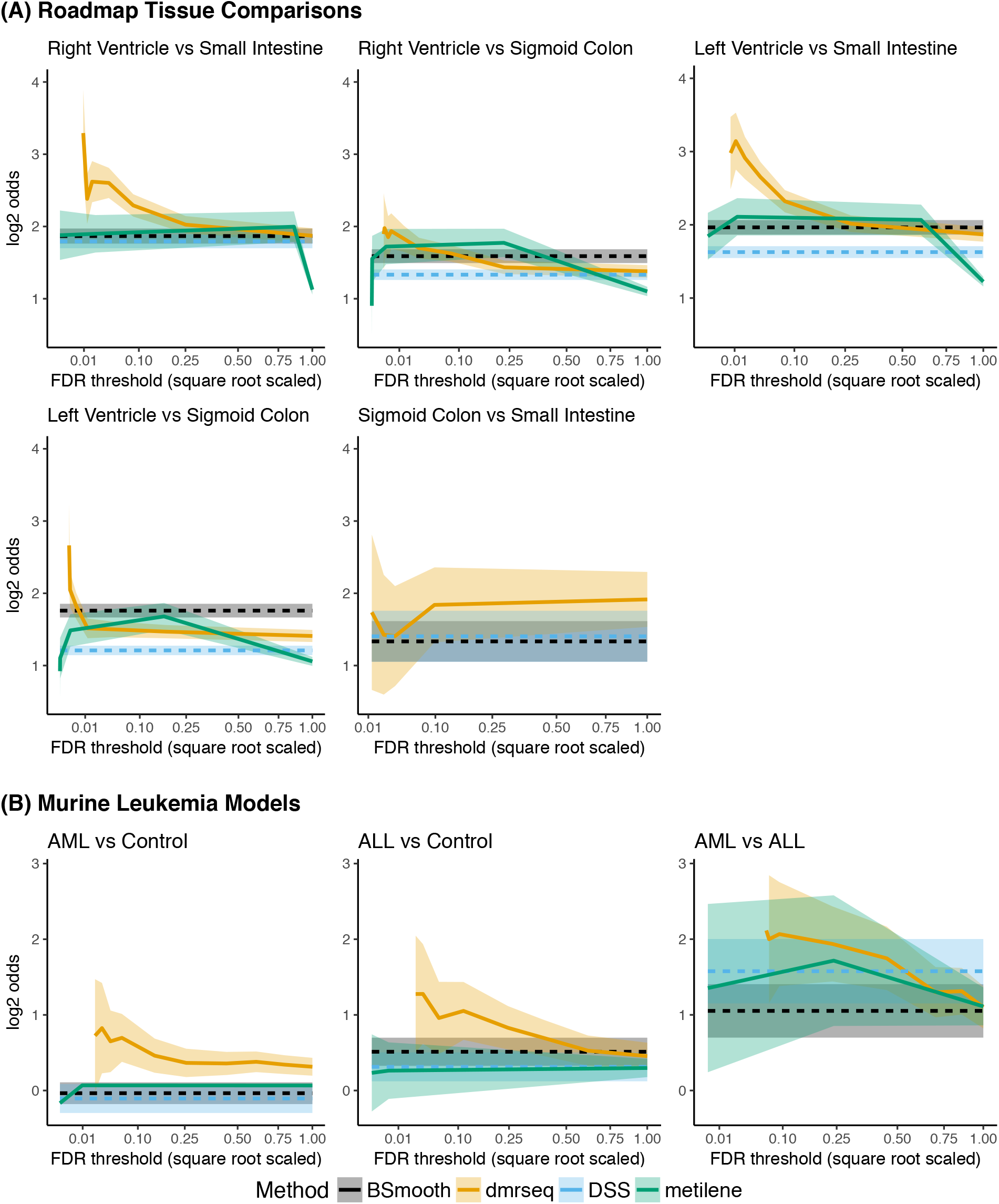
dmrseq achieves stronger inverse association of methylation and differential expression at lower FDR thresholds. Odds of inverse association between methylation difference of (A) tissue-specific DMRs and (B) murine leukemia DMRs with differential expression of nearby DE genes (log2 transformed) is displayed on the y-axis. For dmrseq and metilene, the x-axis represents the FDR threshold (square-root scaled) for which the odds calculation (cumulative) is performed. Since FDR cannot be specified for BSmooth or DSS, the odds are calculated over all DMRs identified and displayed as a horizontal line. Note that the comparison between Left and Right Ventricles is not shown, since no DE genes were identified.

#### 4.2.2 DNMT3a loss in murine leukemia models

In the murine leukemia models, dmrseq finds the most DMRs in the comparison of AML and the control (Table 5), which is also the comparison for which the most DE genes were identified (see Supplementary Table S4). In contrast, DSS and metline both find the most DMRs in the comparison with the fewest DE genes identified, and BSmooth identified similar numbers of DMRs in each comparison, each with far more DMRs than the other methods.

**Table 5:** Murine Leukemia model DMR results. Number of DMRs found by dmrseq and metilene at FDR level 0.10, and BSmooth and DSS under default settings.

| Condition Comparison | dmrseq | BSmooth | DSS | metilene |
| --- | --- | --- | --- | --- |
| AML vs Control | 16,465 | 43,818 | 21,256 | 3,004 |
| ALL vs Control | 8,855 | 51,723 | 16,478 | 3,182 |
| AML vs ALL | 9,253 | 50,004 | 23,582 | 3,360 |

The murine leukemia DMRs found by dmrseq are enriched for inverse associations with DE genes, and this enrichment is stronger for DMRs with lower FDRs (Figure 4). Additionally, enrichment is generally stronger than that of BSmooth, DSS, and metline. While metiline also provides an FDR estimate, there is no consistent association between the FDR ranking and strength of association with expression. Similar to the tissue specificity analysis, BSmooth and DSS DMRs with highest effect sizes exhibit comparable enrichment to dmrseq, with metilene considerably lower, and the enrichment when including all DMRs often drops lower for BSmooth, DSS, or metline than for dmrseq (Supplementary Figures S8 and S9).

## 5. Discussion

We have described dmrseq, a method useful for discovering and prioritizing DMRs from WGBS data. The approach is based on rigorous statistical reasoning and is the first method that permits accurate inference on DMRs that are found by scanning the genome. By developing a transformation that results in summary statistics from candidate regions being exchangeable, we are able to borrow strength across the genome to build a null distribution that permits inference with a sample size as small as 2. We have demonstrated how the method clearly outperforms currently used tools with several experimental data examples and Monte Carlo simulation. The method is implemented as open source software in the form of an R package.

## Supplementary Material

The reader is referred to the online Supplementary Materials for further details of data acquisition and processing, additional methodological details, software implementation details, and supplementary results. In addition, annotated R scripts for the simulation and case study analyses are available in the GitHub repository https://github.com/kdkorthauer/dmrseqPaper, and the R package dmrseq is available on GitHub at https://github.com/kdkorthauer/dmrseq.

## Acknowledgement

Conflict of Interest: None declared

## Funding

The work of all authors was partially by NIH R01 grants HG005220 and GM083084.

